# Zoonotic and Avian Pathogen Detections in Fecal and Sediment Samples – A Low-risk, High-throughput One Health Approach to Surveillance

**DOI:** 10.64898/2025.12.19.694637

**Authors:** Gabriela J. Rzeszutek, Jordan Wight, Mustafa S. Jafri, Abigail J. Erwin, Morgan Hiebert, Ryan Harrigan, Megan Halbrook, Nicole A. Hoff, Isaac Bogoch, Anne W. Rimoin, Jason Kindrachuk, Hannah L. Wallace

## Abstract

Many pathogens, both those with human spillover potential as well as avian-specific viruses, are maintained in wild bird populations. While routine surveillance for influenza A viruses (IAVs) is performed annually, surveillance for other pathogens is limited. Sampling of wild birds is time-consuming, labour-intensive, often limited in sample size, and involves handling of wild and potentially infected birds, posing an increased risk of direct exposure for personnel. Additional methods for surveillance are needed given these significant challenges. Longitudinal fecal and sediment sampling was performed at various sites in southern Manitoba, Canada, particularly focused in Winnipeg from May to October 2025. Sites were chosen based on the suitability of the area for waterfowl habitat, the presence of waterfowl in the area, as well as proximity to reported outbreaks of H5N1 influenza virus. Fecal and sediment samples were collected and screened for the presence of influenza A virus (IAV), Newcastle disease virus (NDV), avian reovirus (ARV), and avian poxvirus (APXV). In total, 782 combined fecal and sediment samples were collected. Of the 714 fecal samples, 34 tested positive for IAV RNA (4.8% prevalence). None of the IAV-positive fecal samples tested positive for H5 RNA. Of the 68 sediments, 15 were positive for IAV RNA (22.1% prevalence), four of which were positive for H5 RNA. NDV RNA positivity was low, with only four positive fecal samples (0.6% prevalence) that were all collected on the same day. ARV RNA positivity was also low, with five positive sediment samples (7.4% prevalence in sediment samples). None of the samples tested positive for APXV DNA. This study builds on previous work showing the utility of environmental sampling for a variety of avian and zoonotic pathogens using a One Health approach that is low-risk, efficient, and high-throughput.

## 1.0 Introduction

Zoonotic viruses have been responsible for the significant portion of the major infectious disease outbreaks and public health emergencies of international concern over the last three decades, including the 2003-2004 severe acute respiratory syndrome epidemic, 2009 influenza A (H1N1), the 2014-2016 West African Ebola virus disease epidemic, coronavirus disease 2019 (COVID-19), and mpox in 2022 and 2024 [1]. The emergence of highly pathogenic avian influenza (HPAI) H5N1 among farmed geese in southern China and subsequent identification of infections among humans in Hong Kong in 1997, coupled with high mortality, raised concerns regarding the epidemic and pandemic potential for this virus. Ongoing circulation of H5N1 among wild birds has resulted in recurrent, unpredictable outbreaks among poultry species, with infrequent infections among humans. The introduction of HPAI H5N1 into North America in 2021 has caused substantial outbreaks among wild birds, domestic poultry, many mammalian species, including cattle, and some human infections [2–9]. These have resulted in increasing concern regarding the ongoing evolution of the virus and its potential to evolve capacity for sustained mammal-to-mammal or human-to-human transmission, as well as considerable economic and food security considerations given ongoing agricultural impacts [10–13].

Despite the risk posed by zoonotic viruses, animal surveillance and sampling has been limited, including in North America. Wild birds, specifically waterfowl, are the one of the major vectors of pathogens that can significantly impact wild bird populations, the agriculture industry, and humans in the context of spillovers [14–16]. While surveillance for influenza A viruses (IAVs) occurs on an annual basis, this work is time-consuming, labour-intensive, involves physically handling and manipulation of wild birds, and thus is often limited in sample size. Additionally, the handling of potentially infected birds poses an increased risk of direct exposure for personnel. Therefore, easier, higher-throughput, lower-effort sampling approaches that mitigate risk are needed to further enhance surveillance.

The use of fecal and sediment samples for surveillance allows greater numbers of samples to be collected, while decreasing specialized training required to perform the sampling, increasing the speed of sample collection, and removing direct interactions with wild birds. Since many of these viruses, including IAVs, are detectable in fecal samples and in the environment for up to a year later [17,18], these sample types are ideal candidates for environmental surveillance.

Although prior studies have reported that detection sensitivities may be lower for environmental samples as opposed to oral and cloacal swabs [19–21], these methods allow for detection of viral pathogens that are present in low viral loads and are particularly useful for large-scale surveillance programs over a wide geographic range. Additionally, given the ability to greatly increase sample sizes and sampling frequencies, environmental sampling allows for pathogens with very low prevalence rates to be detected [22–24], lending additional statistical power to detect accurate prevalence rates.

To demonstrate the utility of environmental surveillance for a variety of zoonotic and avian viruses, this study conducted targeted surveillance for specific viral pathogens based on a number of key considerations. These included pathogens that i) can have a high burden of disease or a high prevalence rate among wild birds; ii) are often found circulating in wild birds and can significantly impact agricultural industries; iii) when present, cause visible signs of infection but may not be particularly virulent; and/or iv) pose zoonotic infection risks for humans.

For this investigation, we selected four viruses as targets for this surveillance effort, each having some or all of the qualities listed above: IAVs, Newcastle Disease virus (NDV, also known as avian paramyxovirus 1), avian reovirus (ARV), and avian poxvirus (APXV). These viruses can all cause significant disease in wild and/or domestic birds and have diverse virology, ecology, pathology, and transmission patterns [25–28].

IAVs are enveloped, segmented, negative-sense RNA viruses from the *Alphainfluenzavirus* genus of the *Orthomyxoviridae* family [25]. IAVs can be classified as low or high pathogenicity viruses with LPAIVs circulating regularly in waterfowl populations, usually without overt signs of disease [29,30]. Conversely, HPAIVs display increased virulence and are associated with high fatality rates in poultry [31–33]. Furthermore, these viruses can cause fatal infections in wild and domestic mammals, humans, and some wild bird species. NDV is an enveloped, negative-sense, single-stranded RNA virus from the family *Paramyxoviridae*. This virus ranges in virulence from lentogenic (low) to velogenic (high) in chickens, and waterfowl are one of the most predominant reservoirs [34]. Infections can be fatal in poultry [28], and have been documented to cause human infections, although this is a rare occurrence [35,36]. Avian reoviruses (ARVs) are non-enveloped viruses with segmented, double-stranded RNA genomes from the *Orthoreovirus* genus of the *Reoviridae* family of viruses [27,37], while avian poxviruses (APXVs) are enveloped, double-stranded DNA viruses from the family *Poxviridae* [26]. Both ARVs and APXVs are generally restricted to infecting only birds, can cause visible lesions and significant inflammation, and can result in high fatality rates, particularly among poultry [38,39].

The current work leverages longitudinal low-risk, high-throughput fecal and sediment sampling for surveillance of four zoonotic and avian pathogens in Manitoba, Canada, using a One Health approach.

## 2.0 Methods

### 2.1 Locations and Timepoints

Sampling was performed longitudinally approximately bi-weekly between 23 May 2025 and 29 August 2025, with one additional sampling timepoint in both September and October 2025. Sporadic sampling was also performed in areas near known outbreaks of HPAI in poultry and in areas further from central Winnipeg. Locations were chosen based on known waterfowl habitat and proximity to waterbodies suitable for waterfowl habitat in addition to recently reported H5N1 outbreaks at poultry facilities. Sampling was performed at five sites in Winnipeg: Assiniboine Park (49.873299°, −97.231154°), St. Vital Park (49.829032°, −97.143637°), Kildonan Park (49.944391°, −97.100024°), Lindenwoods (49.832143°, −97.190863°), and Waverley Heights (49.805287°, −97.165046°). Sampling was also conducted in two additional locations in Manitoba, Niverville (49.595145°, −97.029708°) and Wallace (49.937711°, −101.147637°) (**Figure 1**).

**Figure 1.**
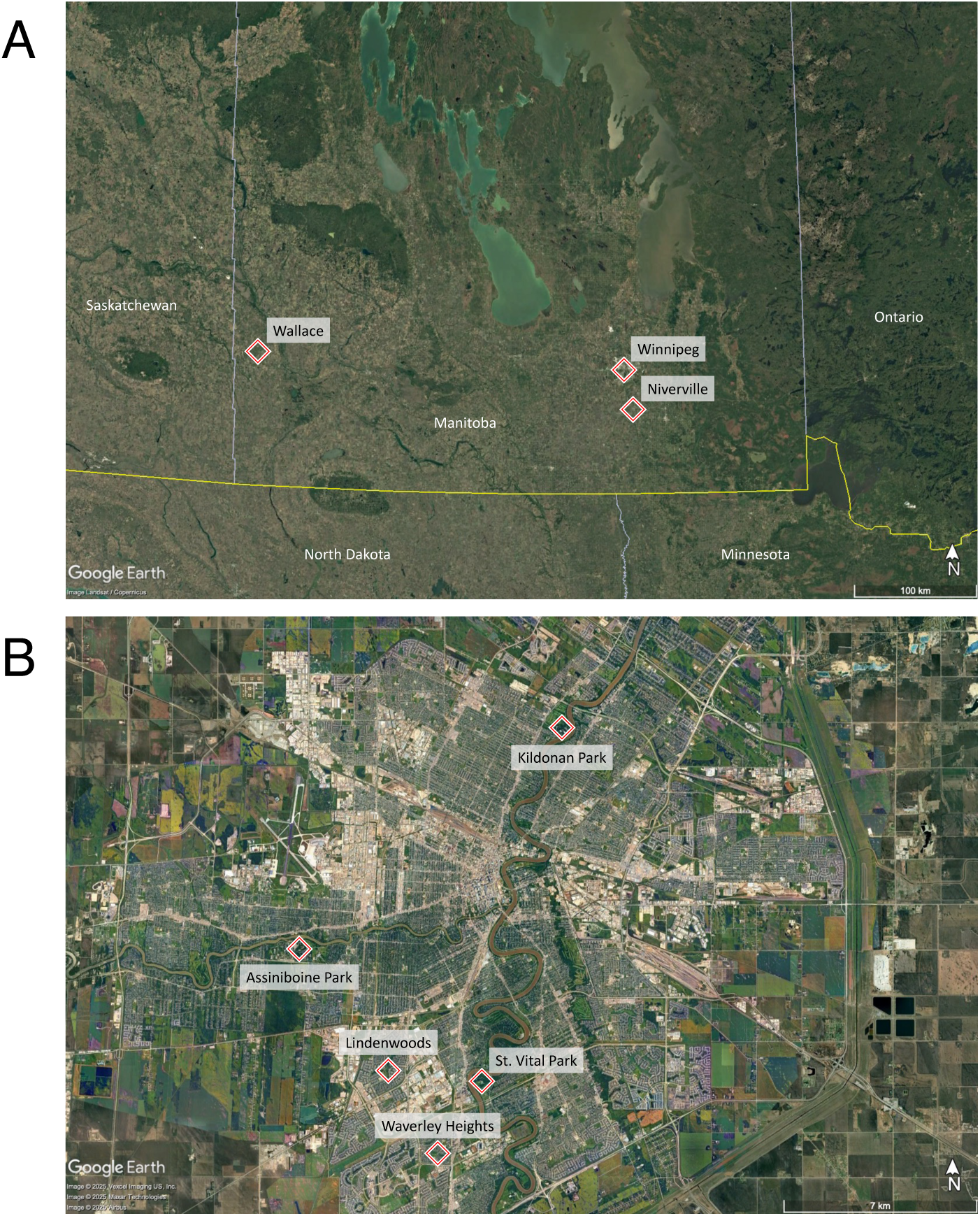
Sampling locations across southern Manitoba, Canada. (**A**) The three municipalities where sampling took place: Winnipeg, Niverville, and Wallace, Manitoba. (**B**) The five sampling locations within the city of Winnipeg: Assiniboine Park, Kildonan Park, St. Vital Park, Lindenwoods, and Waverley Heights.

### 2.2 Fecal Sampling

Fresh fecal samples (either observed being deposited or still visibly wet) from waterfowl, including predominantly Canada Geese (*Branta canadensis*) as well as other dabbling ducks present at the time of sampling, were swabbed and placed in viral transport media (VTM; Puritan™ UniTranz-RT Transport Systems, Fisher Scientific, 22-027-105). Samples were kept on ice until arrival at the lab, at which time samples were vortexed well and aliquoted before being stored at –80°C.

### 2.3 Sediment Sampling

Superficial sediment samples were collected from multiple locations at each sampling site. Sediments were collected in sterile 50mL conical tubes and kept on ice. After arrival at the lab, sediment samples were stored at −80°C until nucleic acid extraction.

### 2.4 RNA Extraction from Swabs

RNA was extraction from 140*μ*L of VTM using the Qiagen Viral RNA Mini Kit (Qiagen, 52906) as per manufacturer’s guidelines. Isolated RNA was stored at −80°C until use.

### 2.5 DNA Extraction from Swabs

DNA was extracted from 200*μ*L of VTM using the DNeasy® Blood & Tissue Kit (Qiagen, 69506) as per manufacturer’s guidelines and stored at −80°C until use.

### 2.6 Nucleic Acid Extraction from Sediments

RNA was extracted from 1g of sediment using the RNeasy® PowerSoil® Total RNA kit (Qiagen, 12866-25) as per manufacturer’s instructions. The RNeasy® PowerSoil® DNA Elution kit (Qiagen, 12867-25) was used to obtain DNA from the same samples following RNA extraction as per manufacturer’s instructions.

### 2.7 Real-time RT-PCR

Real-time RT-PCR was performed using a QuantStudio^TM^ 6 Flex Real-Time PCR System (Applied Biosystems) and analyzed using QuantStudio^TM^ Real-Time PCR Software v1.7.2 (Applied Biosystems). Real-time RT-PCR was performed using AgPath-ID^TM^ One-Step RT-PCR reagents (Applied Biosystems, 4387424). Primer and probe sequences for each of the assays below are included in the supplementary materials (**Supplementary Table 1**). All probes were double-quenched and synthesized by IDT Inc., Canada.

#### 2.7.1 Screening for IAV and H5-specific RNA

Real-time RT-PCR was used to screen RNA for the presence of the IAV matrix gene as per Wight *et al*., 2024 [7]. Any IAV matrix-positive samples were subsequently tested for the presence of the H5 gene as per Wight *et al*., 2024 [7]. Both of these assays are described in detail in [reference 7].

#### 2.7.2 NDV Screening

RNA samples were screened for the presence of the NDV matrix gene by real-time RT-PCR. Screening was performed using 25*μ*L reactions, containing 12.5*μ*L of 2X RT-PCR Buffer, 1.0*μ*L of RT-PCR enzyme mix, 1.0*μ*L of AgPath^TM^ Detection Enhancer (Applied Biosystems, A44941), 0.5*μ*L of 20*μ*M APMV-1-4100F, 0.5*μ*L of 20*μ*M APMV-1-4220R, 0.5*μ*L of 6*μ*M APMV-1-4169P, 4*μ*L of nuclease-free water, and 5*μ*L of RNA. Primers and probes were taken from Wise *et al*., 2004 [34]. The cycling was performed in standard mode as follows: 45°C for 20 min, 95°C for 10 min, 45 cycles of 94°C for 10 sec, 52°C for 40 sec at which time fluorescent signal was detected, followed by 72°C for 10 sec.

#### 2.7.3 ARV Screening

Screening for the M1 gene of ARV was performed in 25*μ*L reactions, containing 12.5*μ*L of 2X RT-PCR Buffer, 1.0*μ*L of RT-PCR enzyme mix, 1.0*μ*L of AgPath^TM^ Detection Enhancer (Applied Biosystems, A44941), 1.25*μ*L of 20 *μ*M ARV_M1_For, 1.25*μ*L of 20*μ*M ARV_M1_Rev, 1.25*μ*L of 10*μ*M ARV_M1_Probe, 1.75*μ*L of nuclease-free water and 5*μ*L of RNA. Primers and probes were taken from Tang and Lu, 2016 [40] with modifications to the reverse primer. Cycling was performed in standard mode as follows: 45°C for 20 min, 95°C for 10 min, and 45 cycles of 94°C for 10 sec and 60°C for 1 min at which time fluorescent signal was detected.

### 2.8 Conventional PCR Screening for APXV

PCRs to screen for the DNA polymerase gene (*pol*) of APXVs were performed using DreamTaq^TM^ Green PCR Master Mix (ThermoFisher Scientific, K1081). Each 25*μ*L reaction volume was comprised of 12.5*μ*L of 2X DreamTaq^TM^ Master Mix, 0.5*μ*L of 10*μ*M PPolF, 0.5*μ*L of 10*μ*M PPolR, 6.5*μ*L of nuclease-free water, and 5*μ*L of DNA. Primers were taken from Gyuranecz *et al*., 2013 [26]. PCR cycling was as follows: 95°C for 5 min, 35 cycles of 94°C for 30 sec, 50°C for 30 sec, and 72°C for 75 sec, with a final extension at 72°C for 7 min. PCR products were subjected to electrophoresis on a 1.5% agarose gel, and DNA bands were visualized using SYBR^TM^ Safe DNA Gel Stain (ThermoFisher Scientific, S33102), imaged using a Gel Doc^TM^ EZ Imaging System (BioRad), and analyzed using ImageLab v6.1 (BioRad).

### 2.9 Positive Controls

Positive controls included RNA isolated from cell-culture propagated human-origin IAVs (A/Macha/01453/2021(H1N1); for IAV matrix screening) and RNA isolated from paired oral/cloacal swabs from an H5 RNA-positive bird from Newfoundland and Labrador, Canada (gift from A.S. Lang, Memorial University; for H5 screening; A/Northern_Fulmar/NL/FAV-0375-2/2023(H5N1)). A synthetic sequence containing the target regions for the NDV, ARV, and APXV assays was synthesized and cloned into pUC57-simple (GenScript) and used as a control for these assays. The synthetic control sequence is included in the supplementary data file.

### 2.10 Maps and Data Visualization

Maps were generated using Google Earth Pro. Data was visualized using R v4.5.0 [41]. R packages included ggplot v3.5.2 [42], and cowplot v1.2.0 [43].

## 3.0 Results

### 3.1 Overall Findings

In total, 714 fecal (**Table 1**) and 68 sediment (**Table 2**) samples were collected between May and October 2025 in and around Winnipeg, Manitoba, for a total of 782 samples. Sampling was dependent on the presence of birds in the area, and therefore sample number varied between sites and between dates. Routine sampling began initially in May of 2025 at three sites, Assiniboine park, St. Vital Park, and Kildonan Park. Opportunistic sampling in the early part of the season was performed in Niverville and Wallace, MB. Wallace was selected in response to detections of HPAI H5N1 in poultry in the area [44]. Later in the season, in response to bird movements, sampling began in Lindenwoods and Waverly Heights (**Table 1**; **Table 2; Supplementary Data**).

**Table 1.** Summary of results for IAV matrix, H5, NDV, ARV, and APXV screening from fecal samples.

| Location | Number of Samples | Number of IAV Matrix Positive Samples | Number of H5 Positive Samples | Number of NDV Positive Samples | Number of ARV Positive Samples | Number of APXV Positive Samples |
| --- | --- | --- | --- | --- | --- | --- |
| Niverville | 28 | 0 | 0 | 0 | 0 | 0 |
| Assiniboine Park | 174 | 1 | 0 | 0 | 0 | 0 |
| St. Vital Park | 181 | 6 | 0 | 1 | 0 | 0 |
| Kildonan Park | 61 | 16 | 0 | 3 | 0 | 0 |
| Lindenwoods | 137 | 0 | 0 | 0 | 0 | 0 |
| Waverley Heights | 133 | 11 | 0 | 0 | 0 | 0 |
| Combined Total | 714 | 34 | 0 | 4 | 0 | 0 |
| % Positivity |  | 4.63% | 0% | 0.54% | 0% | 0% |

**Table 2.** Summary of results for IAV matrix, H5, NDV, ARV, and APXV screening from sediment samples.

| Location | Number of Samples | Number of IAV Matrix Positive Samples | Number of H5 Positive Samples | Number of NDV Positive Samples | Number of ARV Positive Samples | Number of APXV Positive Samples |
| --- | --- | --- | --- | --- | --- | --- |
| Niverville | 15 | 3 | 3 | 0 | 0 | 0 |
| Assiniboine Park | 14 | 3 | 0 | 0 | 0 | 0 |
| St. Vital Park | 18 | 3 | 0 | 0 | 3 | 0 |
| Kildonan Park | 7 | 5 | 1 | 0 | 1 | 0 |
| Lindenwoods | 3 | 1 | 0 | 0 | 1 | 0 |
| Waverley Heights | 2 | 0 | 0 | 0 | 0 | 0 |
| Wallace | 9 | 0 | 0 | 0 | 0 | 0 |
| Combined Total | 68 | 15 | 4 | 0 | 5 | 0 |
| % Positivity |  | 22.06% | 5.88% | 0% | 7.35% | 0% |

### 3.2 IAV Matrix

Of the 714 fecal samples tested, 34 were positive for IAV matrix RNA (4.8%, 34/714 fecal samples). Of the 68 sediment samples tested, 15 were positive for IAV matrix RNA (22.1%, 15/68 sediment samples). IAV positive fecal samples were collected at four of the six fecal sampling locations (Assiniboine Park, St. Vital Park, Kildonan Park, and Waverley Heights) during at least one sampling timepoint. Five of the seven sediment sampling sites had IAV matrix-positive sediment samples (Niverville, Assiniboine Park, St. Vital Park, Kildonan Park, and Waverley Heights) (**Figures 2, 3; Tables 1, 2; Supplementary Data**).

**Figure 2.**
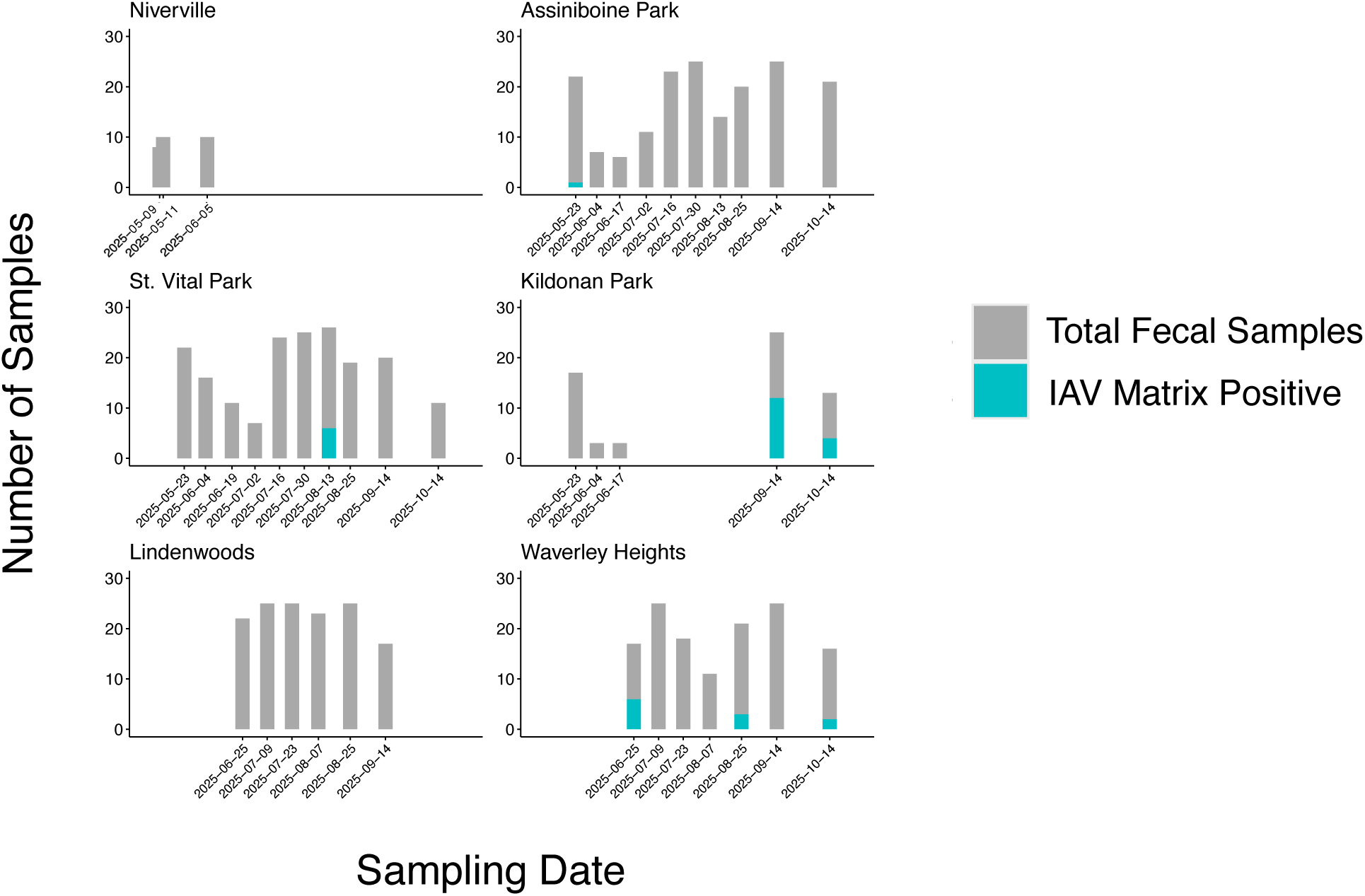
The proportion of IAV matrix positive and negative fecal samples by site and stratified by sampling date. Sampling locations included, Niverville, Assiniboine Park, St. Vital Park, Kildonan Park, Lindenwoods, and Waverley Heights.

**Figure 3.**
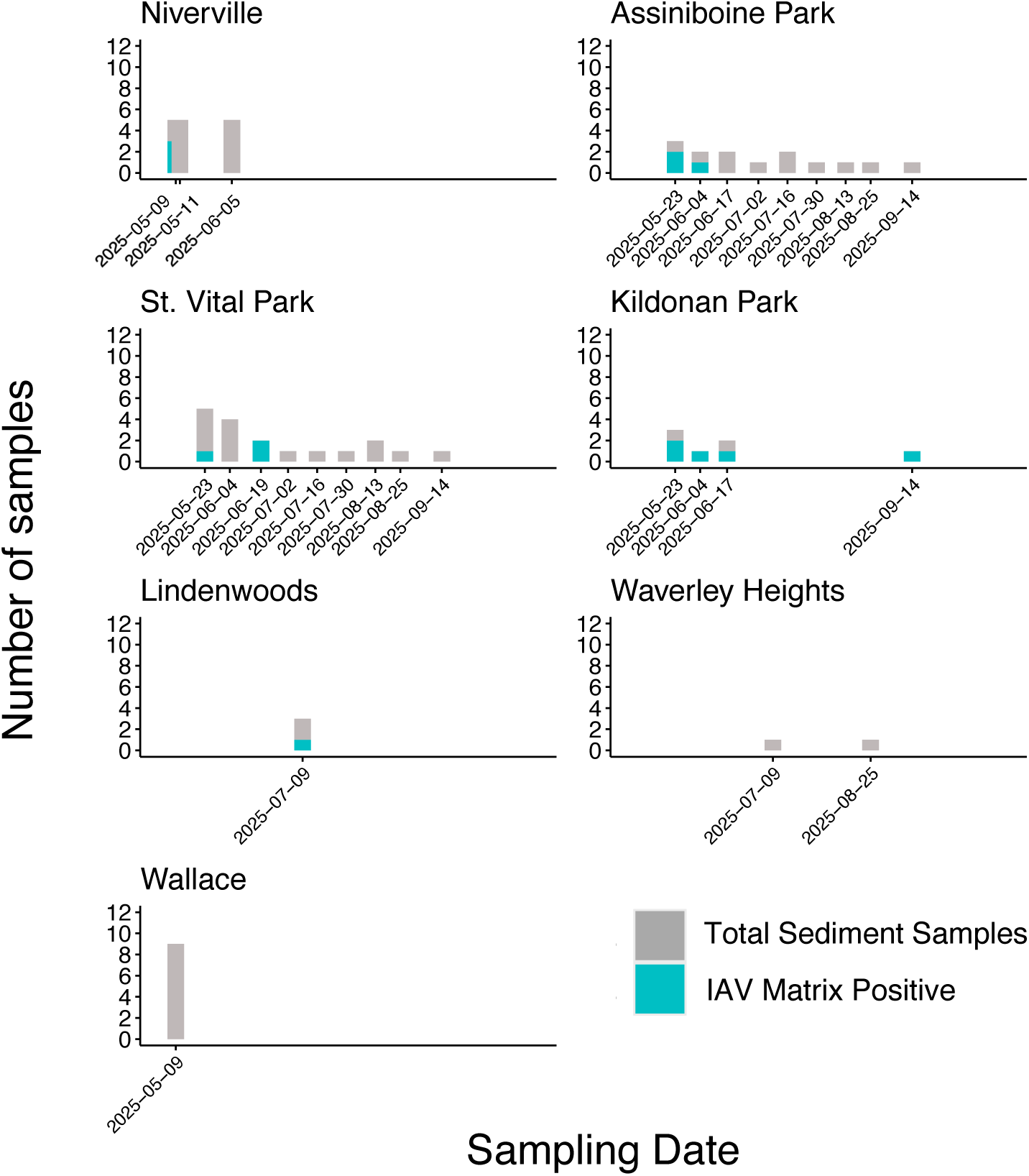
The proportion of IAV matrix positive and negative sediment samples by site and stratified by sampling date. Sampling locations included, Niverville, Assiniboine Park, St. Vital Park, Kildonan Park, Lindenwoods, Waverley Heights, and Wallace.

### 3.3 H5 Screening

Of the 34 IAV matrix-positive fecal samples and 15 of IAV matrix-positive sediment samples, no fecal samples had detectable H5 RNA, while four sediment samples were H5 RNA positive. This equates to a 5.9% prevalence of H5 RNA among all sediment samples (4/68). H5 positive sediments were detected only in Niverville (3/4 H5 positive samples) and at Kildonan Park (1/4 H5 positive samples) (**Tables 1, 2; Supplementary Data**).

### 3.4 NDV Screening

NDV RNA positivity was very low, with four positive fecal samples (0.6%, 4/714) and no sediment samples (0%, 0/68 sediment samples). Interestingly, all four NDV positive samples were obtained on 14 September 2025, with one at St. Vital and three at Kildonan Park (**Figure 4**; **Tables 1, 2; Supplementary Data**).

**Figure 4.**
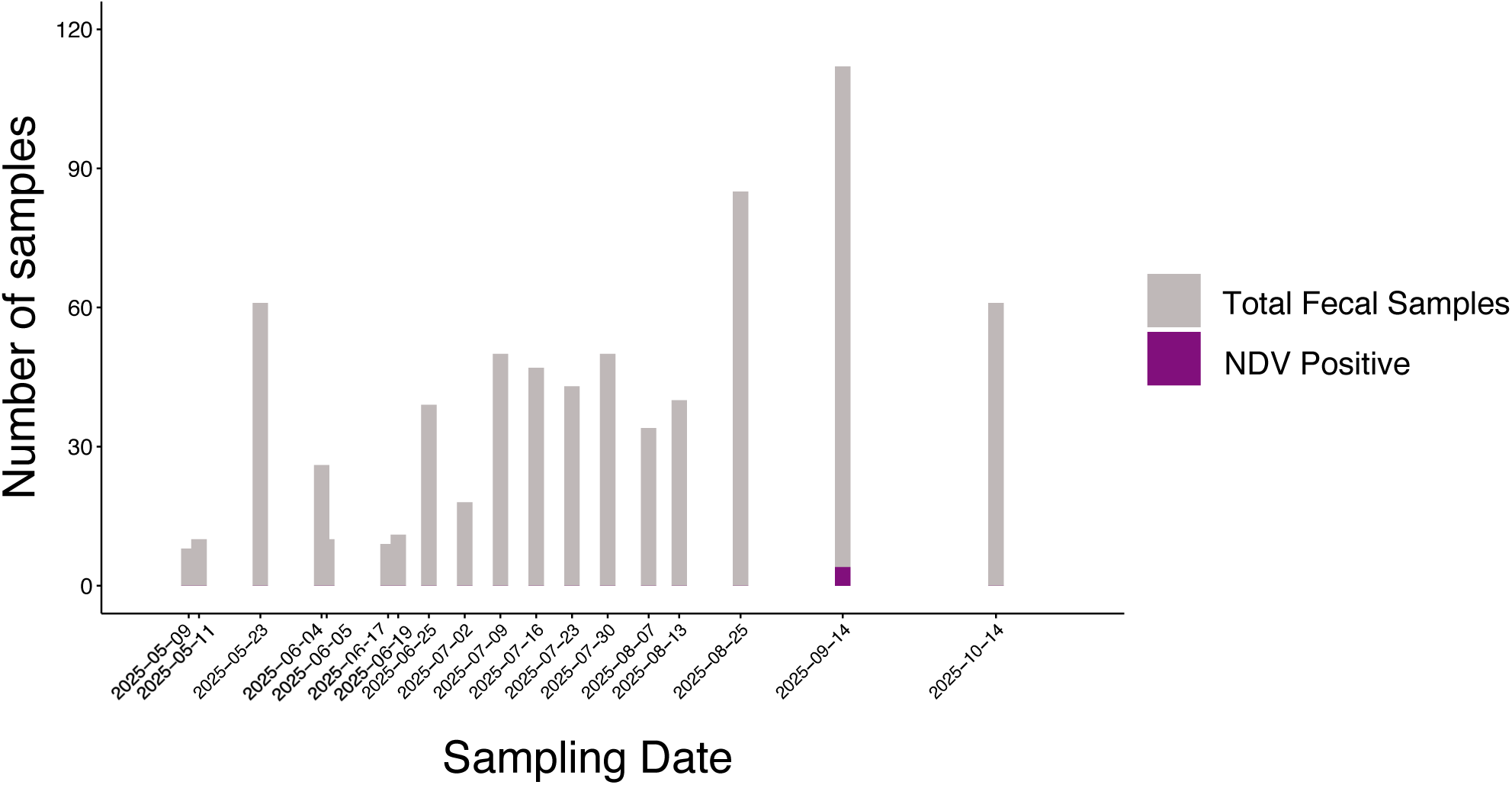
The proportion of NDV positive fecal samples and negative at all locations, stratified by sampling date. Sampling locations included, Niverville, Assiniboine Park, St. Vital Park, Kildonan Park, Lindenwoods, and Waverley Heights.

### 3.5 ARV Screening

ARV positivity was also low, with five of the 68 sediment samples testing positive for ARV RNA (7.4%, 5/68). No fecal samples tested positive for ARV RNA (0%, 0/714). The ARV-positive samples were from three locations, St. Vital Park (3/5 ARV positive samples), Kildonan Park (1/5 ARV positive samples), and Lindenwoods (1/5 ARV positive samples), at various timepoints throughout the study (**Tables 1, 2; Supplementary Data**).

### 3.6 APXV Screening

None of the 782 total samples tested positive for APXV DNA (0%, 0/782). (**Tables 1, 2; Supplementary Data**).

### 3.7 Co-detections

Four of the 782 total samples tested positive for more than one of the pathogens of interest. Two sediment samples tested positive for IAV matrix RNA and ARV RNA, one from St. Vital Park and one from Kildonan Park. Two fecal samples, both collected from Kildonan Park on the same day, tested positive for IAV and NDV RNA (**Supplementary Data**).

## 4.0 Discussion

Given the increasing frequency of zoonotic infectious disease outbreaks [1,45], the recent impacts of these on the health of human, wildlife, and agricultural species, as well as the considerable economic impacts of these outbreaks, the importance of zoonotic pathogen surveillance has never been more clear. Here, we report that environmental surveillance, particularly via fecal and sediment sampling, is highly useful for monitoring IAV prevalence, but also for the detection of other zoonotic and avian pathogens, including NDV and ARV.

Given the labour-intensive nature of live-capture wild bird surveillance, risks associated with the handling of potentially infected animals, required training of staff, and animal use and banding permits, environmental sampling has considerable benefits compared to these traditional approaches. While environmental sampling (fecal and/or sediments) cannot entirely replace live-capture IAV surveillance in waterfowl, it does allow for a supportive supplement to traditional sampling, allowing for increased numbers of samples to be processed without the need to handle the birds directly or have highly-trained personnel to perform sampling.

We detected a 4.8% prevalence rate of IAV in fecal samples. Previous studies have shown that IAV prevalence from swabs is generally higher than in fecal samples [46], suggesting the prevalence rates reported here may be an under-estimate [47]. Since standard IAV surveillance involves the use of paired oropharyngeal and cloacal swabs [7,17,24,46], performing fecal-only surveillance may additionally underestimate the true prevalence of IAVs, particularly due to the lack of sampling the oral cavity (**Figure 2**; **Table 1; Supplementary Data**). While the results presented here may be an underestimate, this approach is sufficiently sensitive to detect IAV RNA in environmental samples and indicates that the true prevalence of IAV among waterfowl is likely higher than was detected among fecal samples in this study.

While the prevalence of IAV was much higher in sediment samples than in fecal samples (**Figure 3**; **Table 2; Supplementary Data**), this was expected. In recent years, it has become clear that although IAVs are enveloped RNA viruses, they are incredibly stable in the environment, with IAV RNA and infectious virus having been shown to persist in the environment for more than a year [17,18]. As a result, interpretation of when virus entered the environment may be difficult, as virus may have been deposited anywhere from moments before sampling to potentially many months prior. Additionally, given the nature of this environmental sampling, we cannot link samples to specific individuals or identify if different samples possibly came from the same bird. Though Canada Geese were highly abundant at the sampled sites and most feces were observed being deposited, other fecal samples may have been from dabbling ducks that were at times present at these sites.

NDVs were detected at very low prevalence, with detections limited to one fecal sample from St. Vital Park and three fecal samples from Kildonan Park (**Figure 4**; **Table 1; Supplementary Data**). This suggests that waterfowl within urban Winnipeg are not frequently infected with NDVs.

ARVs were only detected in sediment samples (**Table 2; Supplementary Data**), which was also as expected. ARVs are non-enveloped viruses, making them highly stable in environmental conditions and are notably resistant to harsh environments, including in sediments and in the gastrointestinal tract of the animals that become infected [38,48].

The environmental sampling performed in this study allowed us to observe site-specific and temporal differences in pathogen detection across southern Manitoba. Of particular interest was that only one sampling site (Kildonan Park) had detections of IAV matrix, H5, NDV, and ARV RNA (**Figures 2, 3; Tables 1, 2; Supplementary Data**). Despite these detections, this site had notably few wild birds present during the middle of the study period, particularly in July and August 2025. This sampling site is more distant from others in urban Winnipeg, being near the northern edge of the city, and is the least influenced by urban developments (**Figure 1B**). We suggest that future studies continue sampling at this site as well as other sites nearby, ideally in combination with a live-capture study of the birds, as a hotspot for the detection of diverse avian pathogens in the environment.

While live-capture IAV sampling and surveillance occurs annually for waterfowl, it is time and resource intensive. This sampling relies upon individuals with a high degree of training who possess necessary permits, as well as the capture and handling of live birds potentially infected with IAVs or other zoonotic pathogens [46]. As such, additional mechanisms for surveillance of not only IAVs, but other avian pathogens are needed. We show here that environmental sampling not only allows for the detection of IAVs, but also other viruses that are often detected at low prevalence, such as NDVs and ARVs. Furthermore, this approach is highly amenable to the addition of surveillance for other pathogens of interest or to return to already collected samples for additional screening.

Through conducting surveillance for zoonotic and avian pathogens in fecal and sediment samples from seven sites across Manitoba, we detected IAV, NDV, and ARV RNA, including at low prevalence. This highlights how environmental sampling can be used to support and supplement ongoing One Health research and how these surveillance efforts are able to detect diverse zoonotic and avian viruses in a manner that is sensitive, high-throughput, and involves low-risk of exposure during sample collection, an approach which may also allow screening for emerging biologic threats.

## Supplementary Materials

**Supplementary Data**. Sample list with screening results.

## Data Availability

Sample list with positivity is included in the **Supplementary Data**. The raw data supporting the findings presented here are available from the authors upon reasonable request.

## Author Contributions

Conceptualization: JW, HLW, JK

Data Curation: GJR, HLW, MH

Formal Analysis: GJR, JW, HLW

Funding Acquisition: JK

Investigation: GJR, MSJ, JW, AJE, MH, HLW

Methodology: JW, HLW

Supervision: JK, HLW

Writing – Original Draft: HLW, GJR

Writing – Review & Editing: All Authors

## Acknowledgements

Sampling assistance was provided by M. Kindrachuk and K. Hiebert. Dr. A.S. Lang (Memorial University) provided H5-positive RNA that was used as a positive control.

## Author Disclosure Statement

This work was done outside of JW and JK’s duties/roles with the Public Health Agency of Canada. The authors declare that they have no other competing interests.

## Funding

This work was supported by the Canadian Institutes of Health Research (Tier 2 Canada Research Chair, grant number 950 231498 to JK) and by the Natural Sciences and Engineering Research Council Discovery Grant (RGPIN-2018-06036 to JK).

---

**Supplementary Table 1.**
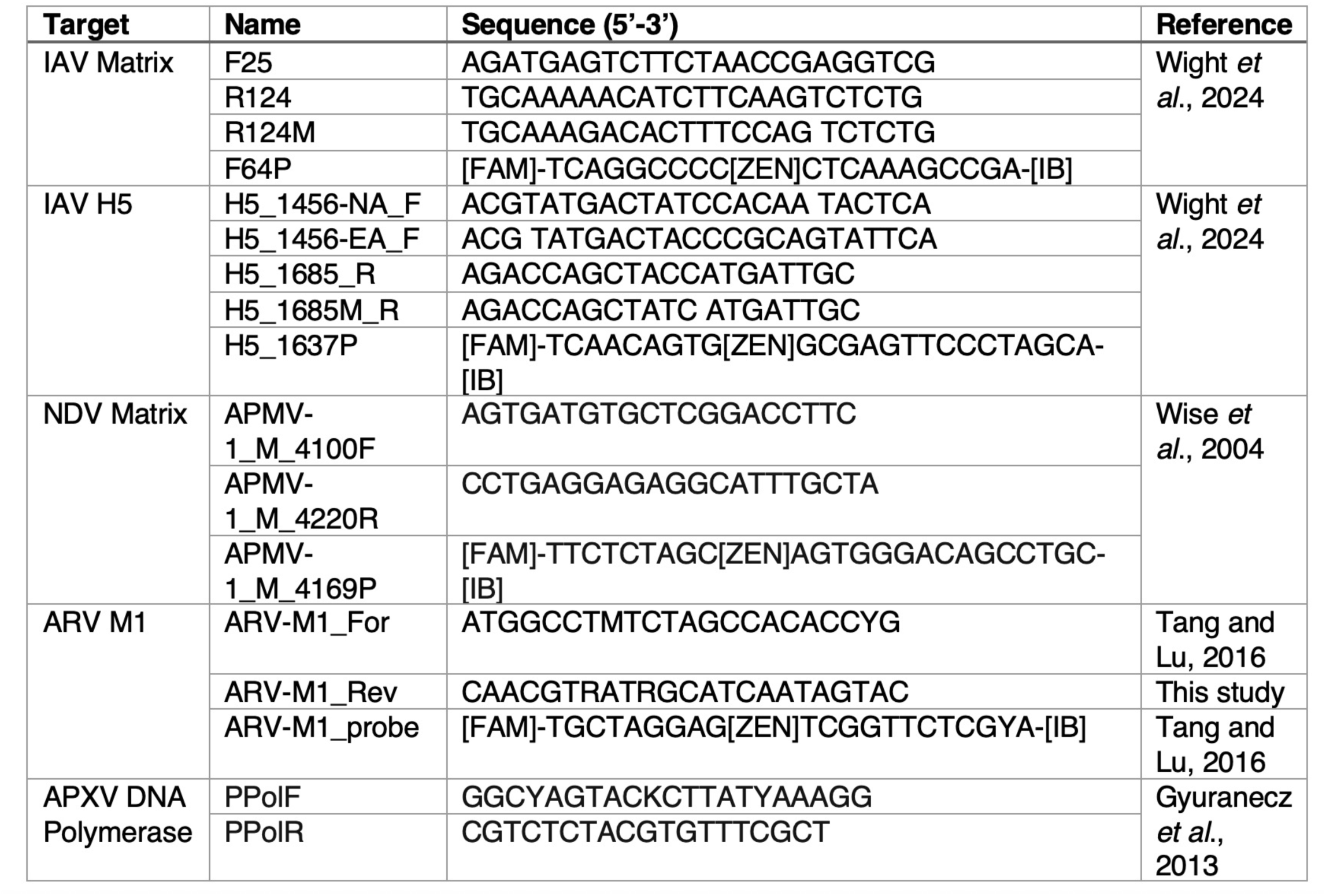
Primer and probe sequences for real-time RT-PCR and conventional PCR screening of fecal and sediment samples.

## Notes

### Competing Interest Statement

The authors have declared no competing interest.

### Summary of Updates

The revised manuscript contains updated numbers relating to screening samples for avian poxvirus.

